# TRPM4 conductances in thalamic reticular nucleus neurons generate persistent firing during slow oscillations

**DOI:** 10.1101/2020.02.11.943746

**Authors:** John J. O’Malley, Frederik Seibt, Jeannie Chin, Michael Beierlein

**Author notes:** Address for Correspondence: Michael Beierlein, Department of Neurobiology & Anatomy, McGovern Medical School, 6431 Fannin Street, Suite 7.046, Houston, TX 77030.

## Abstract

During sleep, neurons in the thalamic reticular nucleus (TRN) participate in distinct types of oscillatory activity. While the reciprocal synaptic circuits between TRN and sensory relay nuclei are known to underlie the generation of sleep spindles, the mechanisms regulating slow (<1 Hz) forms of thalamic oscillations are not well understood. Under *in vitro* conditions, TRN neurons can generate slow oscillations in a cell-intrinsic manner, with postsynaptic Group 1 metabotropic glutamate receptor (mGluR) activation leading to the generation of plateau potentials mediated by both T-type Ca^2+^ currents and Ca^2+^ -activated nonselective cation currents (I_CAN_). However, the identity of I_CAN_ and the possible contribution of thalamic circuits to slow rhythmic activity remain unclear. Using thalamic slices derived from adult mice of either sex, we recorded slow forms of rhythmic activity in TRN neurons, which were mediated by fast glutamatergic thalamoreticular inputs but did not require postsynaptic mGluR activation. For a significant fraction of TRN neurons, synaptic inputs or brief depolarizing current steps led to long-lasting plateau potentials and persistent firing (PF), and in turn, resulted in sustained synaptic inhibition in postsynaptic relay neurons of the ventrobasal thalamus (VB). Pharmacological approaches indicated that plateau potentials were triggered by Ca^2+^ influx through T-type Ca^2+^ channels and mediated by Ca^2+^ and voltage-dependent transient receptor potential melastatin 4 (TRPM4) channels. Taken together, our results suggest that thalamic circuits can generate slow oscillatory activity, mediated by an interplay of TRN-VB synaptic circuits that generate rhythmicity and TRN cell-intrinsic mechanisms that control PF and oscillation frequency.

**Significance Statement:** Slow forms of thalamocortical rhythmic activity are thought to be essential for memory consolidation during sleep and the efficient removal of potentially toxic metabolites. *In vivo*, thalamic slow oscillations are regulated by strong bidirectional synaptic pathways linking neocortex and thalamus. Therefore, *in vitro* studies in the isolated thalamus can offer important insights about the ability of individual neurons and local circuits to generate different forms of rhythmic activity. We found that circuits formed by GABAergic neurons in the thalamic reticular nucleus (TRN) and glutamatergic relay neurons in the ventrobasal thalamus generated slow oscillatory activity, which was accompanied by persistent firing in TRN neurons. Our results identify both cell-intrinsic and synaptic mechanisms that mediate slow forms of rhythmic activity in thalamic circuits.

## Introduction

Slow (< 1Hz) rhythmic activity in the thalamocortical system is a defining feature of natural sleep and is thought to be essential for the grouping of higher frequency activity such as sleep spindles (Beenhakker and Huguenard, 2009; Lüthi, 2014), network plasticity and memory consolidation (Steriade and Timofeev, 2003; Diekelmann and Born, 2010), and homeostatic processes including the clearance of metabolites (Varga et al., 2016). On a cellular level, slow oscillations are characterized by rapid transitions between periods of sustained depolarizations and persistent firing (Up states) and quiescence (Down states), in both neocortex (Steriade et al., 1993b; Sanchez-Vives and McCormick, 2000) and thalamus (Steriade et al., 1993a; Crunelli and Hughes, 2010). It is well established that neocortex plays an important role in the generation of slow oscillations (Steriade et al., 1993b; Sanchez-Vives and McCormick, 2000; Neske, 2016), with cortical activity leading to the recruitment of thalamic targets via extensive corticothalamic projections (Steriade et al., 1993a; Stroh et al., 2013). However, accumulating evidence shows that rather than being passively entrained by cortical afferents, thalamus can generate distinct types of rhythmic activity which in turn shape cortical activity patterns (Rigas & Castro-Alamancos, 2007; Lemieux et al., 2014; Halassa et al., 2011; David et al., 2013; Fernandez et al., 2018). Thus, rhythms in the thalamocortical system might result from the interplay of multiple distinct oscillators (Crunelli and Hughes, 2010). While there is strong consensus on the mechanisms that underlie slow oscillations in neocortex (Crunelli and Hughes, 2010; Neske, 2016), it is important to gain a better understanding of how thalamic neurons and circuits generate slow forms of rhythmic activity.

*In vitro* studies in the isolated thalamus have shown that neurons in the TRN (Blethyn et al., 2006) and in thalamic relay nuclei (Hughes et al., 2002) can act as cellular pacemakers of slow rhythms, under conditions of sustained activation of postsynaptic Group I mGluRs by either exogenous agonists or high frequency corticothalamic activity. This results in an increase in excitability, due to the closure of a K^+^ leak conductance, enabling low threshold Ca^2+^ currents (I_T_) and Ca^2+^-activated non-selective cation currents (I_CAN_) to generate long-lasting and rhythmic plateau potentials, which in TRN neurons lead to persistent firing (Blethyn et al., 2006). However, the precise role of I_T_ and the molecular mechanisms mediating I_CAN_ are not known (Zylberberg & Strowbridge, 2017; Crunelli et al., 2018). Moreover, these findings do not directly address a potential role of intrathalamic networks. TRN and relay neurons form powerful reciprocal connections (Gentet and Ulrich, 2003; Pinault, 2004; Pita-Almenar et al., 2014) which underlie the generation of sleep spindles (Beenhakker and Huguenard, 2009), but whether these circuits can also generate slower forms of oscillatory activity is not known.

Here we addressed these questions by performing recordings in horizontal slices of somatosensory thalamus of adult mice. Surprisingly, we observed highly robust slow oscillatory activity, which was driven by fast synaptic transmission but did not require mGluR activation. We found that a significant number of TRN neurons displayed synaptically-evoked persistent firing (PF), which could also be evoked by brief depolarizing current steps in the absence of synaptic signaling. PF was triggered by Ca^2+^ influx through T-type Ca^2+^ channels and generated by long-lasting plateau potentials mediated by TRPM4 conductances. Our findings highlight how intrinsic properties of TRN neurons and intrathalamic synaptic circuits interact to generate slow thalamic oscillatory activity.

## Materials and Methods

### Animals

We employed C57BL6/J mice (JAX Laboratories, Stock No: 000664) unless specified otherwise. For some experiments, we used TRPC1,4,5,6 general quadruple knockout mice (Sachdeva et al., 2018) kindly provided by Dr. Michael Zhu or TRPC3 general knockout mice (Hartmann et al., 2008) kindly provided by Dr. Oleh Pochynyuk. To optogenetically activate cholinergic inputs to TRN, we used bacterial artificial chromosome (BAC)-transgenic mice expressing channelrhodopsin (ChR2) under the control of the choline acetyltransferase (ChAT) promoter (ChAT–ChR2–EYFP; JAX Laboratories, Stock No: 014546, Zhao et al., 2011). All animals used in this study were treated following procedures in accordance with National Institutes of Health guidelines and approved by the University of Texas Health Science Center at Houston (UTHealth) animal welfare committee.

### Slice Preparation

Brain slices derived from adult animals (P90-120) were prepared as described previously (Ting et al., 2014). Briefly, male or female mice were anesthetized using isoflurane and perfused using an ice cold N-Methyl-D-Glutamine (NMDG) based solution saturated with 95% O_2_–5% CO_2_ consisting of the following (in mM): 92 NMDG, 2.5 KCl, 1.25 NaH_2_PO_4_, 10 MgSO_4_, 0.5 CaCl_2_, 30 NaHCO_3_, 20 glucose, 20 HEPES, 2 thiouera, 5 Na-ascorbate, and 3 Na-pyruvate. Brains were removed and horizontal slices (300 µm) were cut in ice-cold NMDG-based solution using a VT1200 S vibratome (Leica). Slices were then held in NMDG-based solution maintained at 35°C for ∼12 minutes before being transferred to a modified artificial cerebrospinal fluid (ACSF) held at room temperature, consisting of the following (in mM): 92 NaCl, 2.5 KCl, 1.25 NaH_2_PO_4_, 2 MgSO_4_, 2 CaCl_2_, 30 NaHCO_3_, 25 glucose, 20 HEPES, 2 thiouera, 5 Na-ascorbate, and 3 Na-pyruvate. Slices from younger animals (P13-38) were prepared as described previously (Agmon & Connors, 1991). Briefly, mice were anesthetized using isoflurane and then decapitated. Slices were cut in ice-cold cutting solution saturated with 95% O_2_/5% CO_2_ consisting of the following (in mM): 213.4 sucrose, 2.5 KCl, 1.25 NaH_2_PO_4_, 10 MgSO_4_, 0.5 CaCl_2_, 26 NaHCO_3_, and 11 glucose. Slices were then transferred to ACSF consisting of the following (in mM): 126 NaCl, 2.5 KCl, 1.25 NaH_2_PO_4_, 2 MgCl_2_, 2 CaCl_2_, 26 NaHCO_3_ and 10 glucose, incubated at 35°C for 20 minutes and then held at room temperature until used for experiments.

### Electrophysiological Recordings

Recordings were performed in a chamber perfused with ACSF held at 32-34°C using a Warner Instruments TC-324B in-line heater. Cells were visualized via infrared differential interference contrast (IR-DIC) under an Olympus BX51WI microscope equipped with a Dage-MTI IR-1000 camera. Whole-cell recordings were obtained using glass pipettes with a tip resistance of 3-5 MΩ. For current clamp recordings in TRN, we used a potassium-based internal solution consisting of (in mM): 133 K-Gluconate, 1 KCl, 2 MgCl_2_, 0.16 CaCl_2_, 10 HEPES, 0.5 EGTA, 2 Mg-ATP, and 0.4 Na-GTP (adjusted to 290 mOsm and pH 7.3). For voltage clamp recordings of inhibitory postsynaptic currents in neurons of the ventrobasal nucleus of the thalamus (VB), recording pipettes were filled with a cesium-based internal solution consisting of (in mM): 120 CsMeSO_3_, 1 MgCl_2_, 1 CaCl_2_, 10 CsCl, 10 HEPES, 3 QX-314, 11 EGTA, 2 Mg-ATP, and 0.3 Na-GTP (adjusted to 295 mOsm and pH 7.3). For loose-patch recordings of TRN neuronal firing, glass pipettes (3-5 MΩ) were filled with ACSF, and recordings were obtained in voltage-clamp mode, at seal resistances of 50–200 MΩ. To minimize any influence on the cell membrane potential, the holding voltage was adjusted continually to maintain a holding current near 0 pA (Perkins, 2006). In experiments probing TRN oscillatory activity, glutamine (0.3 mM) was added to the ACSF to prevent a rundown of network activity (Bryant et al., 2009; Pita-Almenar et al., 2014). For experiments examining the mechanisms of PF, the bath contained NBQX. In some experiments we replaced extracellular NaCl with equimolar NMDG and used an internal solution consisting of (in mM): 133 K-Gluconate, 1 KCl, 4 NaCl, 2 MgCl_2_, 0.16 CaCl_2_, 10 HEPES, 0.5 EGTA, 2 Mg-ATP, and 0.4 Na-GTP (adjusted to 290 mOsm and pH 7.3). For optogenetic activation of cholinergic afferents we used 1 ms pulses of blue light generated by an LED light source (UHP-T-450-EP, Prizmatix) delivered through a 60x, 0.9 NA water-immersion objective (Olympus) and centered locally in the TRN.

NBQX, apamin, nimodipine, TTX, 2-Aminoethoxydiphenylborane (2-APB), flufenamic acid (FFA), 9-phenthranol, glibenclamide, 4-Chloro-2-[[2-(2-chlorophenoxy)acetyl]amino]benzoic acid (CBA), JNJ 16259685, MTEP hydrochloride, and atropine were obtained from R&D Systems. ω-conotoxin MVII and TTA-P2 were obtained from Alomone labs. All other chemicals were obtained from Sigma-Aldrich.

### Data acquisition

Recordings were made using a Multiclamp 700B amplifier (Molecular Devices), filtered at 3–10 kHz, and digitized at 20 kHz with a 16-bit analog-to-digital converter (Digidata 1440A; Molecular Devices). Data were acquired using Clampex 10.3 software (Molecular Devices) and analyzed using custom macros written in IGOR Pro (Wavemetrics).

### Experimental design and statistical analysis

TRN neuronal firing evoked by brief (25 ms) current steps was classified into two groups, bursting and persistent firing (PF), based on the duration of the evoked membrane depolarization measured at half-amplitude (see *Results*). Post-hoc K means cluster analysis confirmed that classification into two clusters explained the largest amount of variance. We used a depolarization duration of 200 ms to distinguish between bursting and PF. Due to rundown of PF in whole-cell mode, data acquisition in the TRN was limited to 3 min following establishment of whole-cell configuration. Pharmacological experiments examining the mechanisms underlying PF were carried out by performing recordings in the presence of a given antagonist under study, and comparing results with recordings performed under control conditions. All data in a given group were collected from slices derived from a minimum of 2 animals and reported as mean ± SEM. All statistical analyses were performed with Prism 7 software. To evaluate significant differences in the proportion of PF between different experimental groups we performed a chi square test, and report results in the following format: degrees of freedom in parentheses, followed by the chi-square value and p value. To test for significant differences in the duration of evoked membrane depolarization between groups of neurons, we performed an unpaired *t*-test and report results as: degrees of freedom in parentheses, followed by *t* value and *p* value. Statistical significance was set at p < 0.05 and adjusted for multiple statistical comparisons with a Bonferroni correction to correct for pairwise error and to reduce Type I error.

## RESULTS

### Slow oscillatory activity and persistent firing (PF) in thalamic networks

Following activation of postsynaptic Group I mGluRs, individual neurons in the TRN can generate slow oscillatory responses, reminiscent of thalamic activity during sleep (Blethyn et al., 2006). It remains unclear how thalamic synaptic networks contribute to slow oscillatory activity. To examine this issue, we performed experiments in horizontal thalamic slices derived from adult (> 3 month) mice. We and others have previously employed extracellular or optogenetic synaptic stimulation to trigger transient (<2 s) forms of rhythmic thalamic activity *in vitro* (Huguenard and Prince, 1994; Pita-Almenar et al., 2014). Surprisingly, by performing loose-patch recordings from TRN neurons, we observed spontaneous and highly rhythmic patterns of action potential activity that could last several minutes (n = 47 TRN neurons, n = 17 slices). For 12/47 (26%) of recordings, action potential activity was organized as rhythmic bursting (frequency: 1.9 ± 0.1 Hz, n = 12, Fig. 1A). Interestingly, a second group of recordings (17/47, 36%) revealed rhythmic patterns of sustained action potential activity (> 200 ms duration) we termed persistent firing (PF), characterized by an initial high-frequency burst, followed by a long-lasting train of action potentials (average duration: 964.9±256.4 ms, average firing frequency: 175±23.4 Hz, n = 17, Fig. 1B). For neurons displaying rhythmic PF, oscillation frequency was 0.5 ± 0.1 Hz (n = 17), significantly lower compared to the oscillation frequency of cells showing burst firing (*t*(28) = 6.5, *p* = 10^−16^,Fig. 1D). The remaining recordings (n=18) showed more complex patterns of activity, and included neurons that displayed both rhythmic PF and burst firing (n=4, Fig. 1C). These data indicate that burst firing and PF are prominent firing modes in the TRN and are associated with oscillatory activity at distinct frequencies.

**Figure 1.**
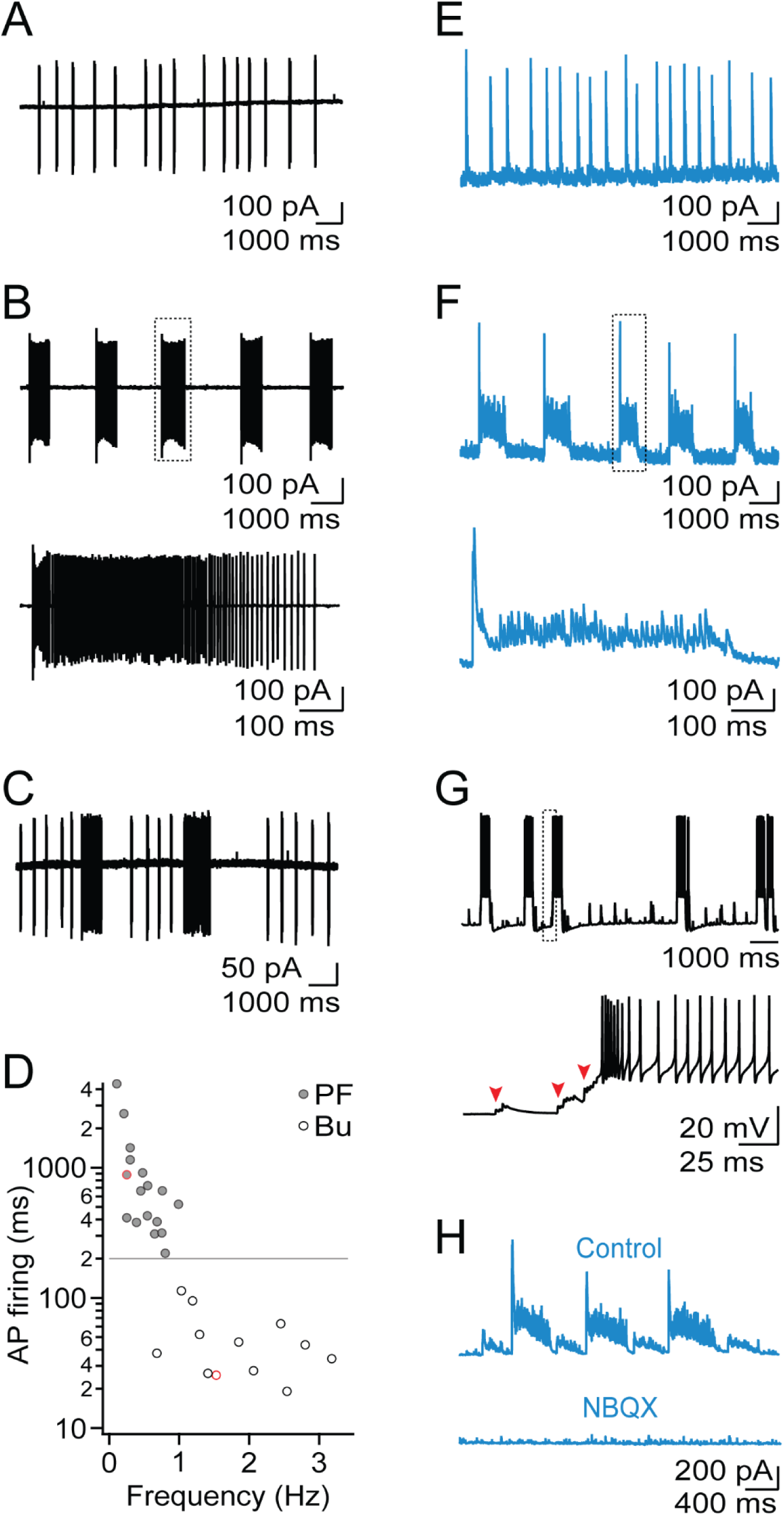
Rhythmic activity in thalamic networks. **A.** Loose-patch recording of TRN neuron showing rhythmic burst firing. **B.** *Top*, TRN neuron showing rhythmic persistent firing (PF). *Bottom*, Close-up highlighting an individual barrage of PF. **C.** TRN neuron showing rhythmic activity consisting of both burst firing and PF. **D.** Summary data (n = 29) plotting the average duration of TRN firing as a function of oscillation frequency, for neurons with either burst firing or PF. Data points in red denote cells shown in A and B. Dashed line marks criterion separating between rhythmic bursting and PF. **E.** Voltage-clamp recording of VB neuron held at 0 mV displaying rhythmic bursts of IPSCs. **F.** *Top*, Recording of VB neuron displaying rhythmic barrages of IPSCs. *Bottom*, close-up shows an individual barrage of IPSCs. **G.** *Top*, Current clamp recording of TRN neuron, showing PF during network activity. *Bottom*, Close up shows that bursts of EPSCs (marked by red arrows) trigger PF. **H.** Representative recording of rhythmic barrages of IPSCs in VB neuron, under control conditions and following bath application of NBQX (10 µM). Similar data were obtained from 4 other VB neurons.

Relay neurons in the ventrobasal nucleus (VB) receive their sole inhibitory input from neurons in the TRN. Therefore, the patterns of synaptic inhibitory responses recorded in VB neurons should closely reflect rhythmic spike firing observed in TRN. To confirm this, we performed voltage-clamp recordings from VB neurons at a holding potential of 0 mV to isolate IPSCs. In 76% (n = 22/29) of all VB neurons that showed oscillatory synaptic activity, we recorded synaptic responses organized as long-lasting high-frequency IPSC barrages (duration: 620.1±54.3 ms, n = 32, Fig. 1F) which occurred rhythmically over several minutes (0.5±0.1 Hz, n = 32). A smaller fraction of recordings showed rhythmic IPSC bursts (n = 4, Fig.1E). Thus, relay neurons are the target of persistent inhibition during slow oscillatory activity, likely mediated by TRN neurons with PF.

Postsynaptic mGluR activation by exogenous agonists or high-frequency stimulation of glutamatergic afferents can lead to slow oscillatory activity in TRN neurons, even in the absence of fast synaptic signaling (Blethyn et al., 2006). In contrast, by performing whole-cell recordings in TRN, we found that rhythmic patterns of PF did not occur spontaneously but instead were evoked by brief bursts of large-amplitude thalamoreticular EPSPs (Fig. 1G). Furthermore, rhythmic IPSCs recorded in VB cells were fully blocked by bath application of NBQX (n = 5, Fig. 1H), suggesting that similar to spindle-like activity (Huguenard and McCormick, 2007), slow oscillatory activity relied on a network of interconnected TRN and relay neurons, at least under our experimental conditions.

Anatomic and functional studies have shown considerable axonal divergence for TRN to VB projections, resulting in highly overlapping TRN afferents to neighboring VB neurons (Pinault, 2004; Pita-Almenar et al., 2014). In agreement, the timing and duration of oscillatory IPSC barrages were highly synchronous for local VB neuronal pairs (< 50 µm somatic distance) recorded simultaneously (Fig. 2A). Synchrony was maintained for individual IPSCs throughout the entire barrage suggesting that the large majority of IPSCs were spike-mediated, without a detectable contribution of asynchronous GABA release (Hefft & Jonas, 2005).

**Figure 2.**
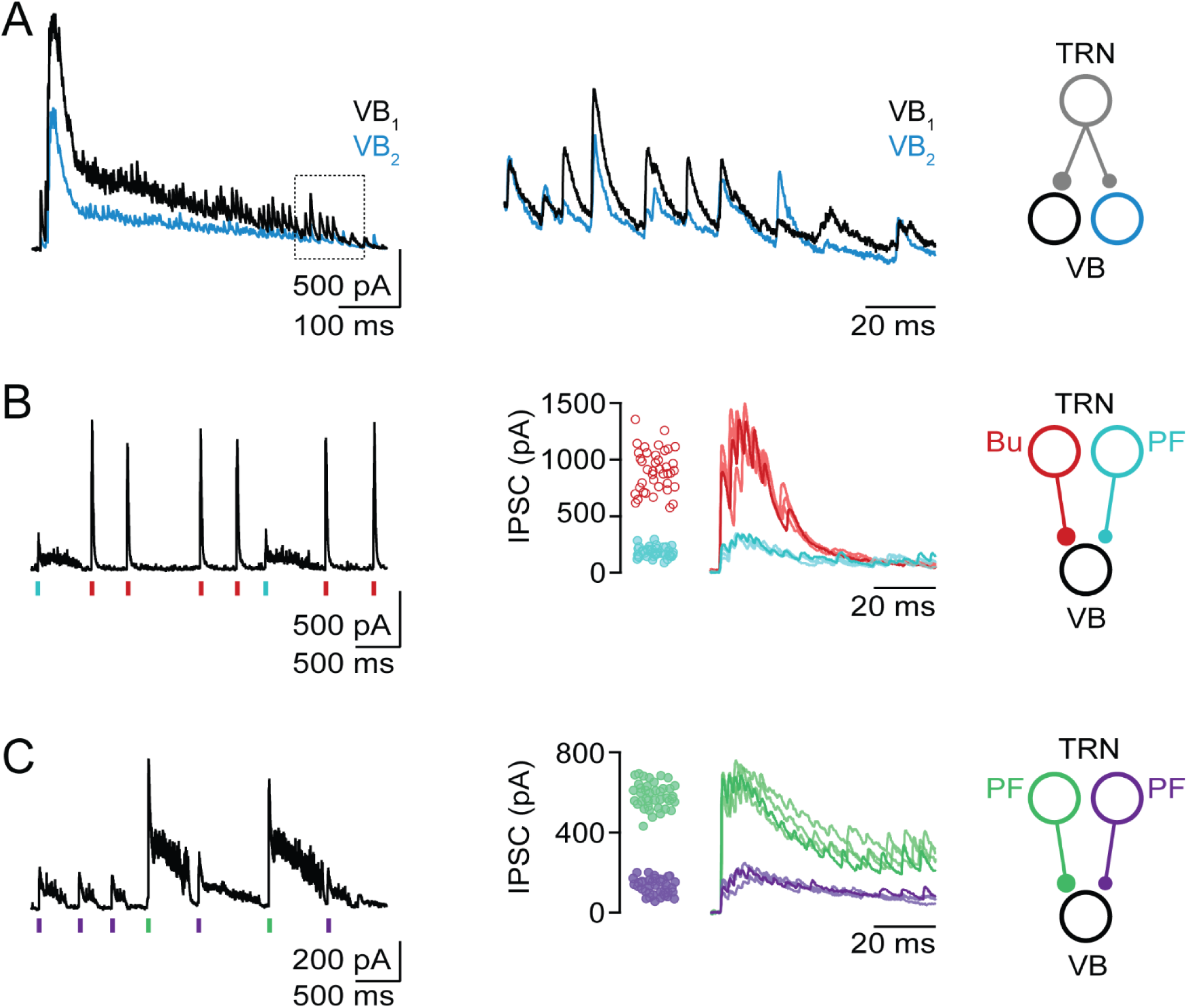
Local TRN neurons display rhythmicity in the absence of precise synchrony. VB recordings were performed in voltage clamp, at a holding potential of 0 mV. **A.** Divergent TRN output to VB neurons. *Left*, Dual recordings of neighboring VB cells showing a single barrage of IPSCs. *Middle*, Close-up of IPSCs (normalized to the initial IPSC amplitude) highlighting precise IPSC synchrony across both cells. *Right*, Schematic outlines proposed divergent TRN projections giving rise to VB responses. **B.** TRN neurons with PF and burst firing form convergent input onto VB neuron. *Left*, Voltage-clamp recording of VB neuron with IPSC bursts (red tick marks) and prolonged IPSC barrages (cyan tick marks) with distinct initial amplitudes. *Middle*, For the same recording, overlay of representative IPSC events and summary data (n=40 for each event) quantifying the initial amplitude of IPSC events. *Right*, Schematic of proposed circuit. **C.** Two TRN neurons with PF form convergent input onto VB neuron. *Left*, Recording of VB neuron showing IPSC barrages with two distinct initial amplitudes (green and purple tick marks, respectively). *Middle*, For the same recording, overlay of representative IPSC events and summary data (n=40 for each event) quantifying the initial amplitude of IPSC events. *Right*, Schematic of proposed circuit.

We found that the majority of VB recordings showed oscillatory IPSC events with little variability of the initial IPSC amplitude and event duration (Fig. 1E,F), consistent with a single presynaptic TRN neuron. However, for a number of recordings we observed either two (n=10 VB cells, Fig. 1H and 2B,C) or three (n=4 VB cells, *data not shown*) different types of rhythmic IPSC events, with each type characterized by a distinct initial IPSC amplitude and event duration, suggesting convergent inputs from two or three presynaptic TRN neurons, respectively. For recordings with two types of events, event duration suggested convergent input by TRN cells that either both displayed PF (n=6, Fig. 2C) or PF and bursting (n=4, Fig. 2B). Importantly, IPSC events evoked by different presynaptic TRN neurons displayed clear rhythmicity but never occurred synchronously. These data confirm our previous findings derived in thalamic slices of less mature animals (Pita-Almenar et al., 2014) and suggest that local thalamic circuits can mediate ongoing oscillatory activity in the absence of widespread synchrony.

### Brief depolarizations trigger PF in TRN neurons

So far our data indicate that a significant fraction of TRN neurons shows PF during network activity. To better characterize the properties and underlying mechanisms of PF, we performed whole-cell recordings and blocked network activity by adding NBQX (10 µM) to the bath solution. Neurons were held at membrane potential of -75 mV unless stated otherwise and actions potentials were evoked using brief depolarizing current steps (25 ms, 400 pA). Such stimuli led to bursts of action potentials (average number of spikes: 8.1±0.2, n = 74) in 67% of neurons recorded (Fig. 3A). For the remaining neurons (37/111, 33%) we observed sustained action potential activity (average number of spikes: 72.2±9.8, firing frequency: 99.1±7.5 Hz, n = 37) closely resembling PF observed during network activity. PF was mediated by long-lasting plateau potentials (amplitude from rest: 29.3±0.8 mV, plateau duration: 807.1±41.7 ms, n = 37, Fig. 3A,B) that terminated in a stepwise manner. Using the duration of the current-step evoked membrane depolarization, we performed K means cluster analysis, confirming that TRN firing patterns can be most readily classified into two distinct groups, bursting and PF, with the latter group displaying membrane depolarizations of > 200 ms. Neurons with current-step evoked PF were not restricted to a specific TRN subregion and were localized along the entire thickness of the TRN shell (not shown). We found that PF remained stable for several minutes but then decayed over time, regardless of composition of the internal solution (Fig. 3C). This might partly explain why studies employing whole-cell methods have not reported PF in adult TRN neurons (e.g. Clemente-Perez et al., 2017), while at least one study using sharp electrode recordings described PF in a subset of neurons of the cat perigeniculate nucleus (Kim and Mccormick, 1998). For the remainder of this study we limited data acquisition to the first 3 minutes in whole-cell mode.

**Figure 3.**
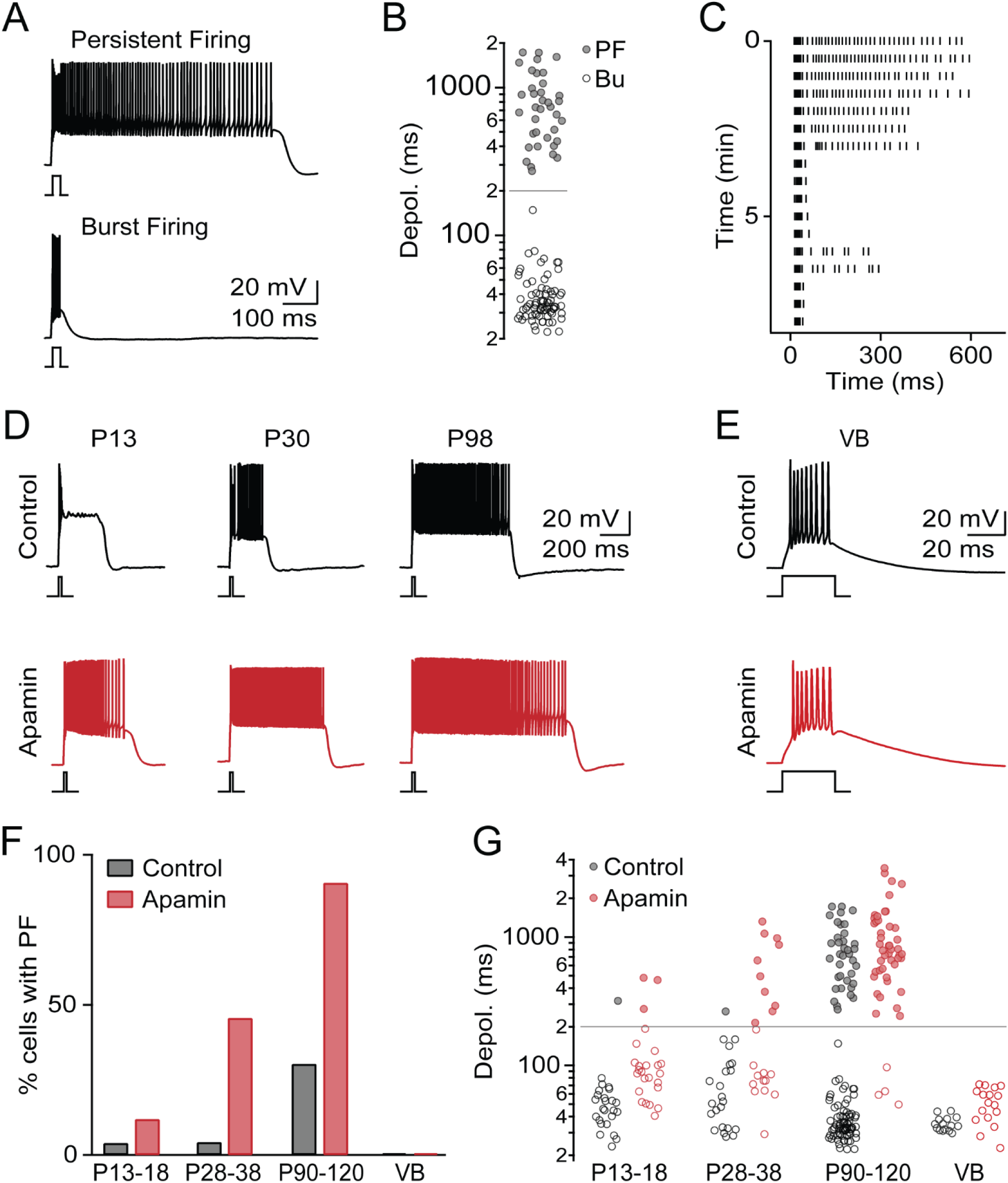
PF occurs late in development and regulated by SK conductances. **A.** Brief depolarizing current steps (25 ms, 400 pA, RMP = -75 mV) lead to two distinct firing patterns, Persistent Firing (PF, *top*) or Burst Firing (Bu, *bottom*). **B.** Summary data (n = 111) showing bimodal distribution of current-step evoked membrane depolarizations. Grey line indicates criterion separating PF and Bu. **C.** Raster plot of AP activity in representative TRN neuron showing rundown of PF. **D.** Representative recordings of TRN neurons with PF at three developmental stages, in control conditions (*black*) and in the presence of apamin (500 nM, *red*). **D.** Representative recordings of TRN neurons with PF at three developmental stages, in control (black) and in the presence of apamin (500 nM, *red*). **E.** Representative recordings of VB cells, in control (black) and in the presence of apamin (red) show lack of PF in both conditions. **F.** Summary data showing incidence of PF in TRN across development (P13-18: n = 24 in control, n = 25 in apamin, P28-38: n = 23 in control, n = 22 in apamin, P90-120: n = 111 in control, n = 50 in apamin) and in VB (n = 15 in control, n = 16 in apamin). **G.** For the same cells as in F, summary data quantifying membrane potential duration for all TRN cells across development, in control conditions and in apamin.

### PF occurs late during development and is controlled by SK conductances

Previous work characterizing TRN firing patterns in slices derived from younger (<P30) animals has not reported PF (Lee et al., 2007), suggesting that PF is developmentally regulated. To examine this possibility, we carried out additional recordings in slices prepared from mice aged P13-18 and P28-38. Compared to adult mice, only 1/24 (4%) of cells at P13-18 and 1/23 (4%) of cells at P28-38 displayed PF (Fig. 3D-F), indicating that PF is prominent only in fully mature animals.

The generation of plateau potentials that underlie PF is likely to be tightly regulated by K^+^ conductances, including small conductance Ca^2+^-activated K^+^ channels (SK). To examine this possibility, we performed recordings in the presence of the SK channel blocker apamin (100 nM). Under these conditions a much larger fraction of TRN cells in adult animals showed PF (92%, n = 46/50 *χ*^*2*^(1) = 47.5, *p* = 10^−16^, Fig. 3D-F). Thus, excitatory conductances responsible for PF are widely expressed in the adult TRN but appear to be strongly controlled by SK conductances. Given the role of SK channels in regulating PF in the adult, the developmental increase in PF we observed might be solely mediated by a progressive decrease of SK expression. However, we found that even in the presence of apamin the incidence of PF remained very low early in development and dramatically increased with animal age (P13-18: 12%, n = 25, P28-38: 41%, n = 24, P90-120: 92%, Fig. 3D-F), suggesting a developmental upregulation of excitatory mechanisms that underlie PF. Finally, we examined if neurons in the VB displayed plateau potentials in control or in the presence of apamin. In contrast to TRN, PF was completely absent from VB under both conditions (Fig. 3E,G). Thus, the mechanisms underlying PF do not appear to be uniformly expressed in all thalamic nuclei.

### Metabotropic glutamate receptor activation is not necessary for PF

Previous work has highlighted a critical role of postsynaptic group I mGluR activation in enabling intrinsic oscillations and Up states in TRN neurons (Blethyn et al., 2006). While PF under our experimental conditions did not require exogenous or synaptic mGluR I activation, it is possible that mGluRs were tonically activated by ambient glutamate (Crabtree et al., 2013) or in a ligand independent manner (Sun et al., 2016). To examine this possibility, we probed TRN firing in the presence of the non-competitive mGluR1 antagonist JNJ 16259685 (JNJ) and the non-competitive mGluR5 antagonist MTEP. Block of group I mGluRs did not significantly change the incidence of PF (control: 33%, n = 111, JNJ+MTEP: 48%, n = 27, *χ*^*2*^(1) = 1.7, *p* = 0.19) or the plateau potential duration for cells with PF (control: 807.1±41.7 ms, JNJ+MTEP: 599.8±64.3 ms, *t*(46) = 1.85, *p* = 0.071, Fig. 4A,B). These data suggest that under our experimental conditions Group I mGluR activation is not essential for PF.

**Figure 4.**
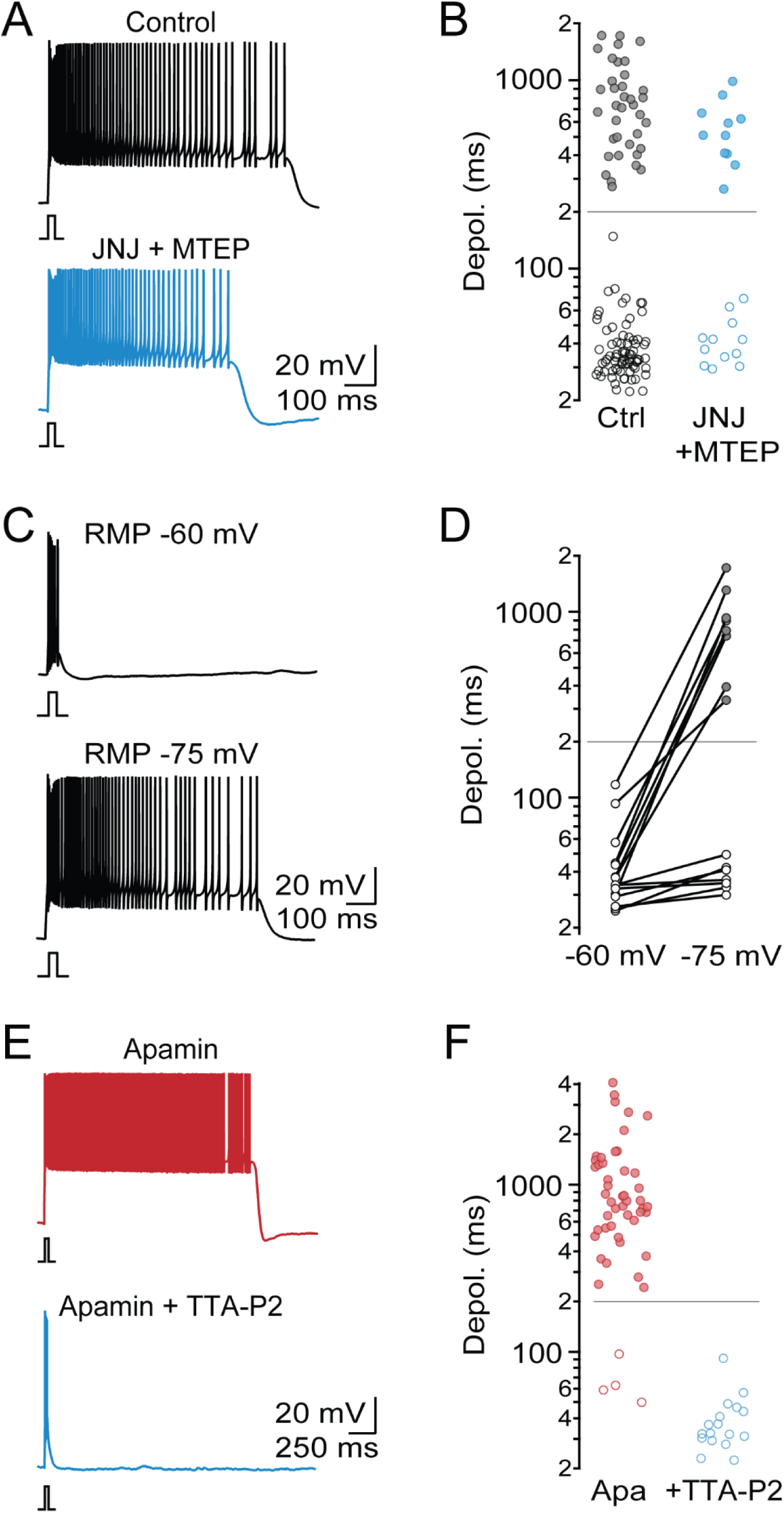
T-type Ca^2+^ channels but not mGluRs are required for PF. **A.** Representative recordings of PF from two different TRN neurons, in control conditions and in the presence of the mGluR1 antagonist JNJ 16259685 (1 µM) and the mGluR5 antagonist MTEP (10 µM). **B.** Summary data quantifying duration of evoked membrane depolarization in control (n = 111) and in the presence of mGluR 1/5 antagonists (n = 23). **C.** Representative recording of neuron with PF, initially held at a holding potential of -60 mV and then at -75 mV. **D.** Summary data quantifying evoked depolarization duration as a function of holding potential (n = 15 TRN neurons). **E.** Representative recordings in the presence of apamin and in the presence of apamin and the T-type Ca^2+^ channel antagonist TTA-P2 (1 µM). **F.** Summary data quantifying evoked depolarization duration (n = 50 in apamin, n = 17 in apamin + TTA-P2).

### Activation of T-type Ca^2+^ channels is required for PF

Next, we examined if PF is voltage-dependent, by evoking action potential activity from two distinct holding potentials. TRN cells held initially at -60 mV never displayed PF or plateau potentials (0/15 cells, Fig. 4C,D). However, when the holding potential was adjusted to -75 mV, 8/15 neurons generated PF (*χ*^*2*^(1) = 1.7, *p* = 0.0013), suggesting that the underlying mechanisms of PF are voltage dependent.

To further isolate the mechanisms responsible for PF, we employed a pharmacological approach. Due to the washout of PF in whole-cell mode, it was not possible to quantify changes in PF prior to and following bath application of antagonists for a given neuron. Therefore, we performed all recordings in the presence of a given antagonist and made comparisons to neurons recorded under control conditions (see *Materials and Methods*). To limit the number of recordings required for meaningful statistical comparisons, we performed the following experiments in the presence of apamin, which, as described above, results in PF in 92% of neurons (Fig. 3E,F).

Previous studies have indicated that in thalamic neurons T-type Ca^2+^ channels are critical for the generation of mGluR-dependent Up states (Hughes et al., 2002; Blethyn et al., 2006). The voltage-dependence of PF we observed appears to be consistent with these results. To confirm the involvement of T-type Ca^2+^ channels in generating PF, we performed recordings in the presence of the specific antagonist TTA-P2 (1 µM). Under these conditions, PF was completely eliminated (apamin: 92%, apamin+TTA-P2: 0%, n = 17, *χ*^*2*^(1) = 46.5, *p* = 10^−20^, Fig. 4E,F). Thus, T-type Ca^2+^ channels are required for the generation of PF.

### The plateau potential underlying PF is mediated by a sodium current

It is possible that long-lasting plateau potentials in TRN neurons are entirely mediated by T-type Ca^2+^ conductances (Zylberberg and Strowbridge, 2017). While these channels show rapid inactivation following strong depolarization, their inactivation is not complete for smaller depolarizations near the resting membrane potential, resulting in a steady-state inward (“window”) current (Williams et al., 1997). However, PF observed in the present study was accompanied by large membrane depolarizations that do not appear to be compatible with a significant role of T-type Ca^2+^ channels. Therefore, we systematically examined the possible involvement of other voltage- or Ca^2+^-gated non-inactivating conductances. We first tested the role of high-threshold Ca^2+^ channels (L, P/Q, and N). In the presence of the selective L-type antagonist nimodipine (nim, 20 µM, Fig. 5A) the incidence of PF (apamin: 92%, apamin+Nim: 80%, n = 10, *χ*^*2*^(1) = 0.3, *p* = 0.6, Fig. 5B) and plateau potential duration for cells with PF (apamin: 1108.5±126.6 ms, apamin+Nim: 846.8±235 ms, *t*(48) = 0.9, *p* = 0.38, Fig. 5C) remained comparable to responses recorded in apamin. Similarly, incubating slices in the P/Q and N antagonist ω-conotoxin MVIIC (ω-Ctx, 1 µM) did not lead to significant changes in the incidence of PF (apamin: 92%, apamin+ω-Ctx: 70%, n = 10, *χ*^*2*^(1) = 1.4, *p* = 0.23, Fig. 5B), and plateau potential duration (apamin: 1108.5±126.6 ms, apamin+ω-Ctx: 726.7±196.2 ms, *t*(47) = 1.2, *p* = 0.24, Fig. 5C), suggesting that high-threshold Ca^2+^ channels do not play a critical role in PF.

**Figure 5.**
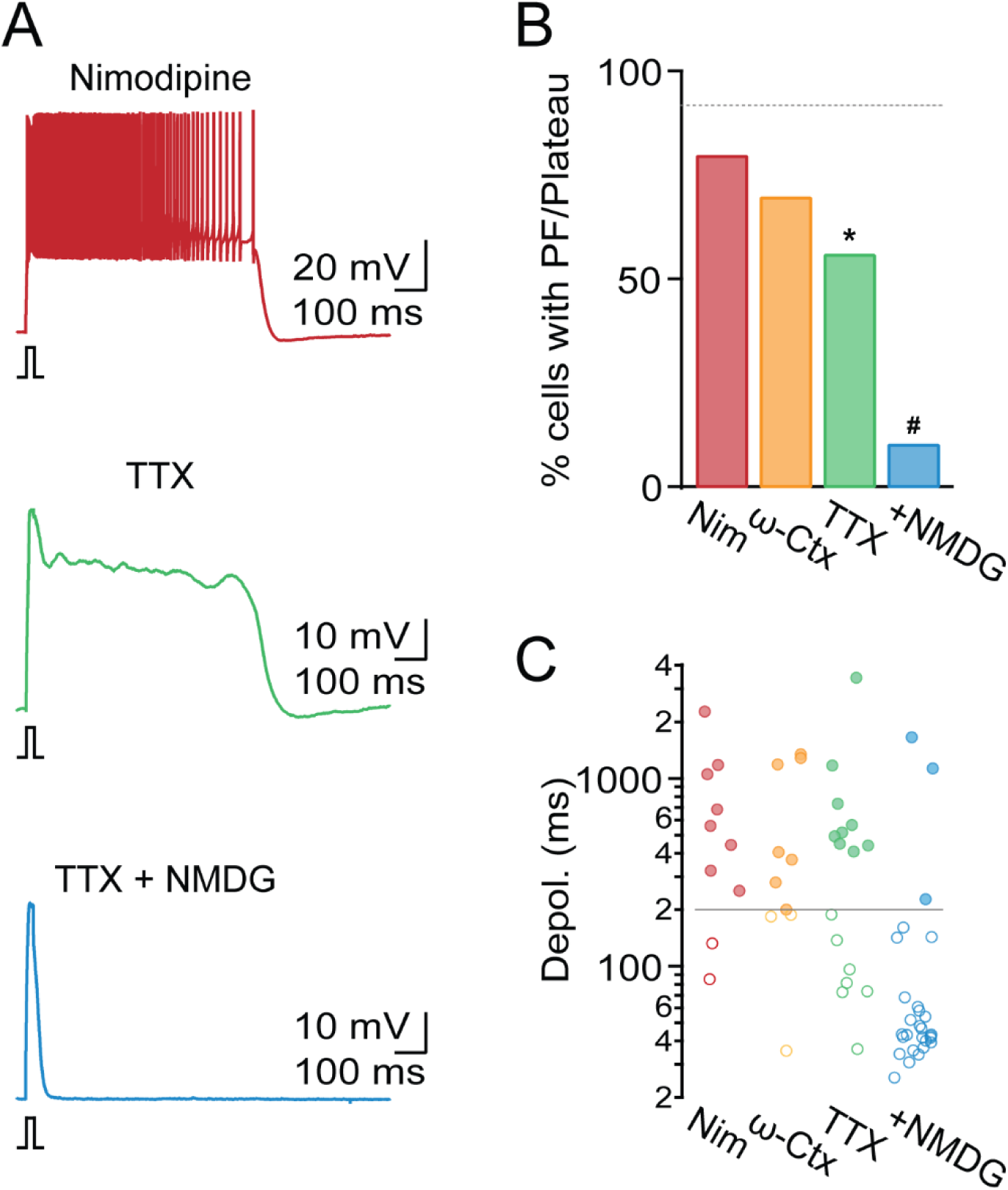
A TTX-insensitive sodium current generates plateau potential. All recordings were performed in the presence of apamin. **A.** Representative TRN recordings in the presence of the L-type Ca^2+^ channel antagonist nimodipine (20 µM, *top*), in TTX (500 nM, *middle*), and in the presence of TTX and NMDG replacing extracellular Na^+^ (*bottom*). **B.** Summary data showing incidence of PF in the presence of nimodipine (Nim, n = 10), the P/Q- and N-type Ca^2+^ channel antagonist ω-conotoxin MVIIC (ω-Ctx, 1 µM, n = 10), TTX (n = 16), and TTX and NMDG (n = 28). Dashed line indicates incidence of PF in apamin. **C.** Summary data from the same cells as in B quantifying evoked depolarization duration in each condition. Grey solid line represents criterion for PF. * denotes statistical significance of p < 0.05 compared to recordings in apamin group, # denotes statistical significance of p < 0.05 compared to recordings in TTX.

It is possible that plateau potentials are mediated by activation of a persistent Na^+^ current (I_NaP_) as was suggested previously (Kim and Mccormick, 1998; Fuentealba et al., 2005). We found that in the presence of apamin and TTX (500 nM), fast action potential activity was completely blocked, revealing a plateau potential in 9/16 cells (amplitude: 30±1.2 mV, n = 16, Fig. 5A), a smaller fraction compared to conditions in apamin (apamin: 92%, apamin+TTX: 56%, *χ*^*2*^(1) =10, *p* = 0.002, Fig. 5B). Plateau potential duration remained comparable to control (apamin: 1108.5±126.6 ms, apamin+TTX: 912.8±325.7 ms, *t*(49) = 0.7, *p* = 0.5, Fig. 5C). These data indicate that voltage-gated Na^+^ currents facilitate plateau potential generation, but are not strictly required.

Next, we examined if Na^+^ currents other than the ones carried by voltage-gated Na^+^ channels are essential for plateau potentials, by substituting the majority of extracellular Na^+^ with NMDG. We found that in the presence of NMDG (together with apamin and TTX) the incidence of plateau potential generation was significantly reduced (apamin + TTX: 56%, apamin + TTX + NMDG: 11%, n = 28, *χ*^*2*^(1) = 10.6, *p* = 0.001, Fig. 5A-C), suggesting that PF is mediated at least in part by a TTX-insensitive Na^+^ current.

### TRPM4 channels underlie PF in the TRN

Our data thus far are consistent with a role of the Ca^2+^-activated nonselective cationic current (I_CAN_) in mediating plateau potentials. Transient receptor potential (TRP) channels have been previously proposed as molecular mechanisms mediating I_CAN_ (Launay et al., 2002), with transient receptor potential canonical (TRPC) and melastatin (TRPM) channels as the most likely candidates. To examine the potential involvement of TRPC channels, we performed recordings in slices derived from global TRPC3 null mutant mice (TRPC3 KO, Hartmann et al., 2008) and from global quadruple TRPC1,4,5,6 null mutant mice (QKO, Tian & Zhu, 2018), in the presence of apamin. For both mouse lines, the incidence of PF (apamin: 92%, TRPC3 KO: 70%, n = 10, *χ*^*2*^(1) = 3.4, *p* = 0.65, QKO: 70%, n = 10, *χ*^*2*^ (1) = 3.4, *p* = 0.65, Fig. 6B) and plateau potential duration (apamin: 1108.5±126.6 ms, TRPC3 KO: 841.1±136 ms, *t*(47) = 0.9, *p* = 0.39, QKO: 752.6±133.4 ms, *t*(47) = 1.1, *p* = 0.27, Fig. 6C) were comparable to our observations in WT animals, suggesting that at least 5 of the 7 TRPCs known to be expressed in the brain are not involved in mediating PF.

**Figure 6.**
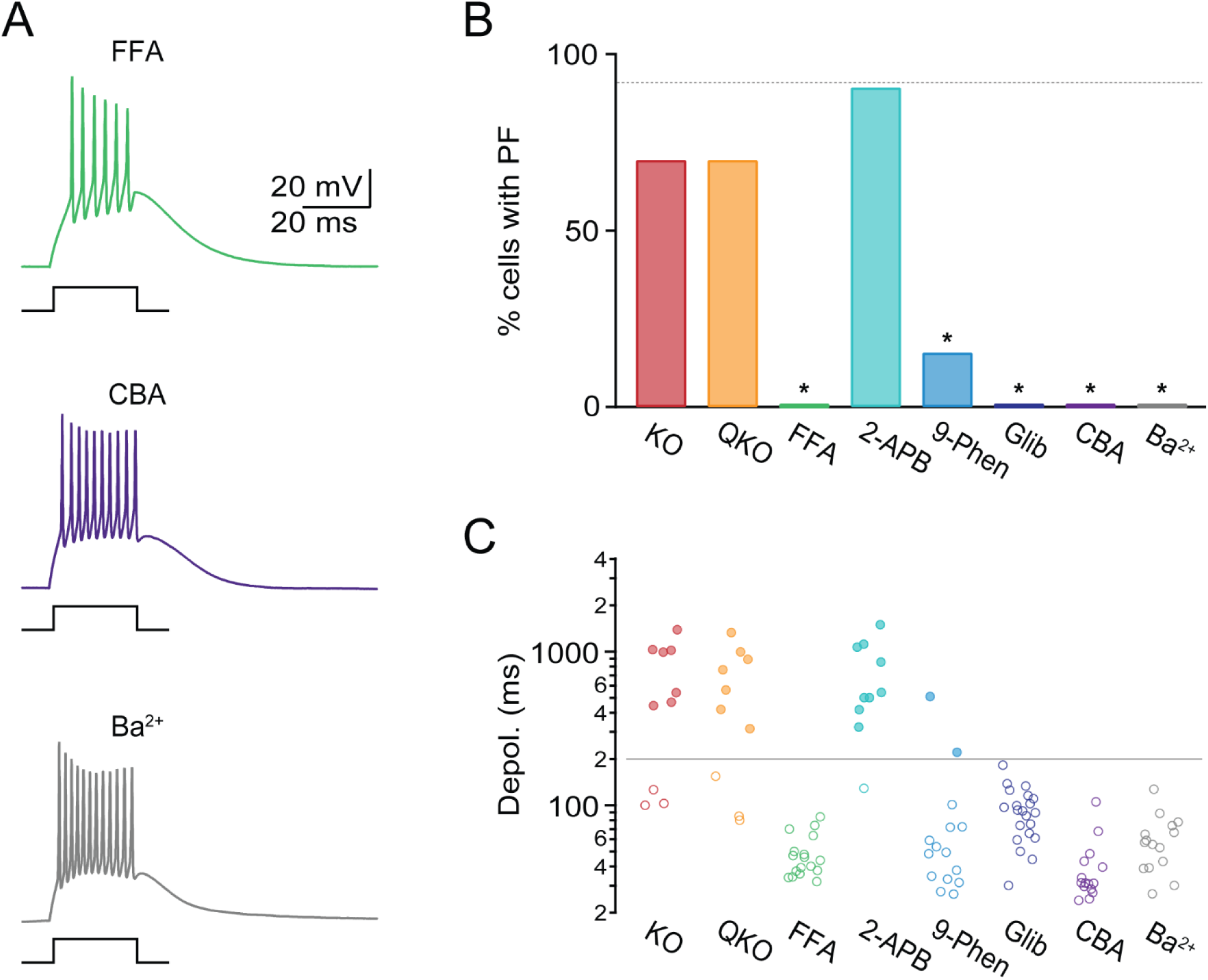
TRPM4 conductances mediate PF. All recordings were performed in the presence of apamin. **A.** Representative recordings in control and in the presence of the non-selective transient potential receptor (TRP) channel antagonist FFA (100 µM, *top*), the TRPM4 antagonist CBA (50 µM, *middle*), and in 2 mM BaCl_2_ replacing extracellular CaCl_2_ (*bottom*). **B.** Summary data showing incidence of PF in TRPC3 KO (KO, n = 10 cells) in TRPC1,4,5,6 quadruple KO (QKO, n = 10 cells), FFA (n = 17), the non-selective TRP channel antagonist 2-APB (50 µM, n = 10), the TRPM4/5 antagonists 9-phenthranol (9-Phen, 100 µM, n = 15) and glibenclamide (Glib, 100 µM, n = 21), CBA (n = 15), and Ba^2+^, (n = 15). Grey dashed line represents incidence of PF in apamin. **C.** For the same cells as in B, summary data quantifying the evoked depolarization duration. * denotes statistical significance of *p* < 0.05 compared to recordings in apamin.

Next, we investigated a possible role of TRPM channels in mediating PF. TRPM2 has been shown to facilitate burst firing (Lee et al., 2013), while both TRPM4 and TRPM5 have been implicated in the generation of plateau potentials in cortex (Lei et al., 2014) and of slow inward currents in cerebellar Purkinje cells (Kim et al., 2013). We found that in the presence of the TRP channel inhibitor FFA (100 µM), PF was completely eliminated (apamin: 92%, apamin+FFA: 0%, n = 17, *χ*^*2*^(1) = 46.6, *p* = 10^−20^, Fig. 6A-C). Next, we examined the effects of the broad-spectrum inhibitor 2-APB which among TRP channels blocks TRPC7 and TRPM2, but does not inhibit TRPM4 or TRPM5. 2-APB had no effect on PF (apamin: 92%, apamin+ 2-APB: 90%, n = 10, *χ*^*2*^(1) = 0.4, *p* = 0.54, Fig. 6B), with plateau potential duration comparable to control conditions (apamin: 1108.5±126.6 ms, apamin+2-APB: 758±132.4 ms, *t*(49) = 1.3, *p* = 0.22, Fig. 6C), ruling out a role of TRPM2 or TRPC7 and suggesting that either TRPM4 or TRPM5 might mediate PF. Both channels are voltage-dependent, monovalent-selective, and require increases in intracellular Ca^2+^ for their activation (Launay et al., 2002; Nilius et al., 2003). We found that in the presence of the TRPM4/5 antagonists 9-Phenthranol (9-Phen, 100 µM) and glibenclamide (Glib, 100 µM) PF was completely eliminated (apamin: 92%, apamin+9-Phen: 15%, n = 15, *χ*^*2*^(1) = 34.2, *p* = 10^−14^, Glib: 0%, n = 21, *χ*^*2*^(1) = 51.4, *p* = 10^−26^, Fig. 6B). Next, we performed experiments using the recently developed selective TRPM4 antagonist CBA (Ozhatil et al., 2018). In the presence of CBA (50 µM), PF was completely eliminated (apamin: 92%, apamin + CBA: 0%, n = 15, *χ*^*2*^(1) = 44, *p* = 10^−17^, Fig. 6A,B). Taken together, our results indicate that TRPM4 conductances mediate the plateau potential underlying PF.

Since increases in intracellular Ca^2+^ are required for TRPM4 activation, exogenous buffers such as BAPTA should eliminate PF. However, due to the rundown of PF, experiments relying on the infusion of Ca^2+^ chelators were not feasible. We therefore probed a possible Ca^2+^ requirement of PF by replacing extracellular Ca^2+^ with equimolar Ba^2+^. Ba^2+^ passes through T-type Ca^2+^ channels and generates large inward currents (Huguenard & Prince, 1992), but unlike Ca^2+^ does not lead to TRPM4 activation (Nilius et al., 2004; Yamaguchi et al., 2014). Under these conditions, brief depolarizing steps still led to prominent burst firing, indicating that T-type Ca^2+^ channel dependent dendritic depolarizations were not affected (Fig. 6A). However, none of the cells displayed PF (apamin: 92%, apamin+Ba^2+^: 0%, n = 15, *χ*^*2*^(1) = 44, *p* = 10^−17^, Fig. 6B,C). These data further confirm that T-type Ca^2+^ channels alone are not sufficient to generate plateau potentials. Instead, our data suggest that T-type Ca^2+^ channel activation and the resulting Ca^2+^ increases lead to the activation of TRPM4 conductances and the generation of PF.

### Synaptic recruitment of muscarinic acetylcholine receptors suppresses PF

Finally, we examined the modulation of PF by synaptic activity. TRN neurons are the target of cholinergic afferents from the basal forebrain and brainstem (Hallanger and Wainer, 1988). Our previous studies have shown that acetylcholine (ACh) release leads to a biphasic postsynaptic response, with fast excitation mediated by nicotinic ACh receptors (nAChRs) and slow and long-lasting inhibition mediated by M2 muscarinic receptors (mAChRs) coupled G protein-gated inward rectifying K^+^ (GIRK) channels (Sun et al., 2013). To examine if cholinergic inputs can trigger PF in the TRN, we performed experiments using BAC-ChAT-ChR2 mice that selectively express ChR2 in cholinergic neurons (Zhao et al., 2011) and optogenetically evoked ACh release with light pulses (1 ms) centered over the recorded neuron. To avoid decay of PF, TRN neurons were recorded in loose-patch mode. Under these conditions, 2/5 of neurons displayed PF (Fig. 7A,C). We reasoned that the activation of mAChRs and the resulting GIRK activation might limit a full manifestation of PF. Indeed, following atropine application to block mAChRs, the duration of action potential firing was significantly prolonged in all neurons recorded (control: 213.5±94 ms, atropine: 416.3±84.1 ms, *t*(4) = 7.1, *p* = 0.002, Fig. 7A-C), and the incidence of PF increased from 40% to 80%. These data show that the activation of K^+^ channels by synaptically released neuromodulators such as ACh can dynamically modulate the expression of PF.

**Figure 7.**
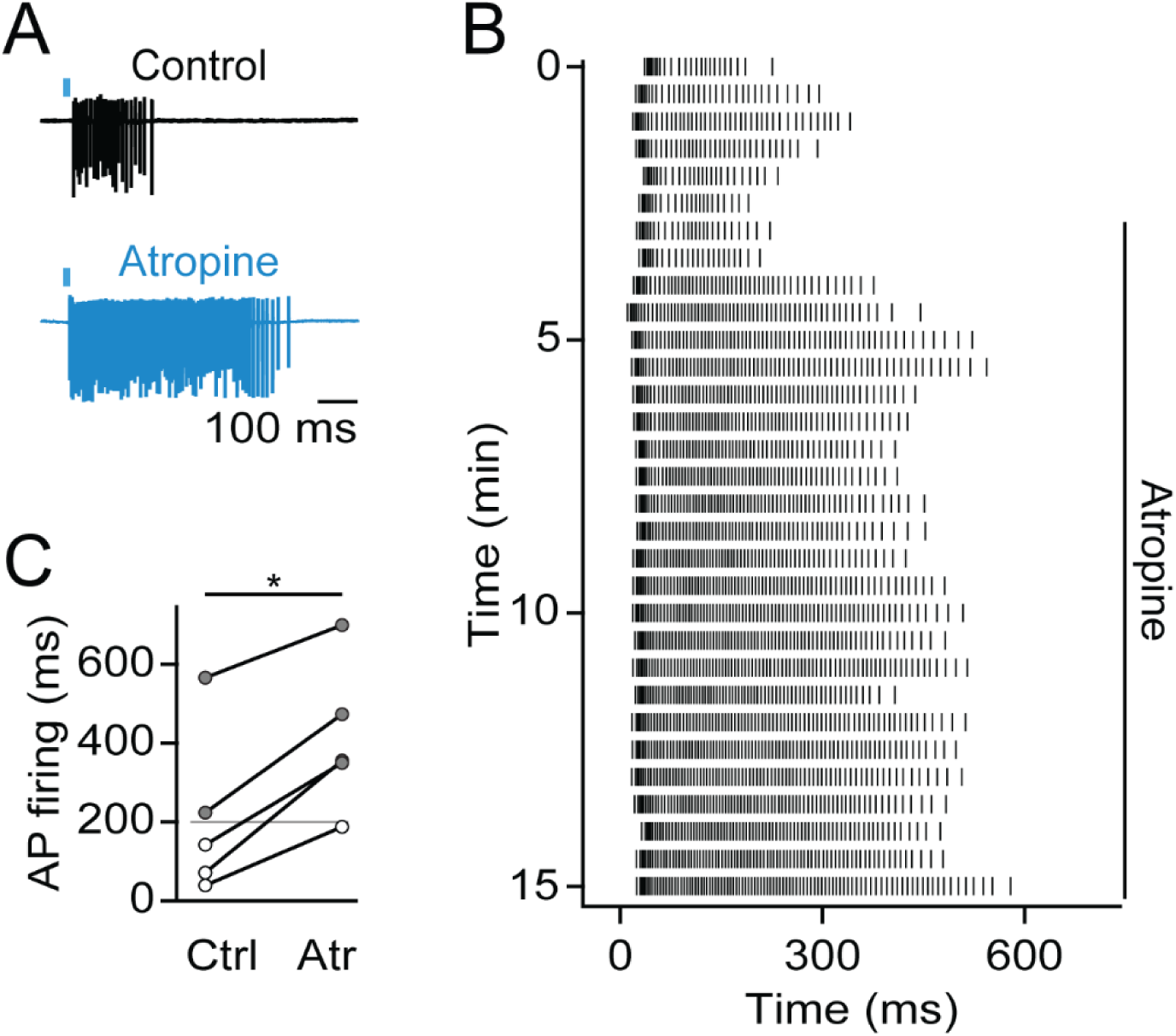
Synaptic recruitment of muscarinic AChRs suppresses PF. Recordings were carried out in slices of ChAT-ChR2-EYFP BAC mice. **A.** Loose-patch recording of a TRN neuron showing brief barrage of ACh-evoked activity (*black*), which is prolonged following application of the mAChR antagonist atropine (10 µM, *blue*). **B.** Raster plot of ACh-evoked neuronal activity prior to and following atropine application, for the cell shown in A. **C.** Summary data quantifying the average duration of ACh-evoked action potential activity before and after atropine application (n = 5). * *p* < 0.05

## Discussion

Here we have identified TRPM4 as the mechanism underlying PF in adult neurons of the TRN. Our data indicate that the large depolarizations and intracellular Ca^2+^ increases required for TRPM4 activation are mediated by T-type Ca^2+^ channels. PF was associated with robust slow oscillatory activity in thalamic circuits which required fast synaptic transmission but did not rely on mGluR receptor signaling. Previous studies have shown that both TRN and thalamic relay neurons can act as cellular pacemakers for slow thalamic oscillations. Our present findings highlight intrathalamic circuits as an alternative mechanism capable of generating slow rhythmic activity.

### Mechanisms of PF in the TRN

Our results extend recent work that has implicated TRPM4 in controlling neuronal excitability in cortex (Lei et al., 2014; Riquelme et al., 2018), hippocampus (Menigoz et al., 2016), substantia nigra (Mrejeru et al., 2011), and brainstem (Picardo et al., 2019). A unique property of TRPM4 activation within physiological ranges is the requirement for both depolarization and large increases in intracellular Ca^2+^ concentration (Launay et al., 2002; Nilius et al., 2003; Nilius et al., 2004). Since dendritic Ca^2+^ signals can experience decay under whole-cell conditions, this might partly explain why we consistently observed rundown of PF in the present study. Alternatively, recordings in whole-cell mode might have led to the dialysis of critical signaling molecules. TRPM4 conductances rapidly desensitize in excised patches (Zhang et al., 2005) and in whole-cell mode, but modulators such as PIP2 can restore TRPM4 activity. Additional investigations into the mechanisms of TRPM4 regulation under physiological conditions will be important.

What are the Ca^2+^ sources of TRPM4 activation? We found that pharmacological block of T-type Ca^2+^ channel using the specific antagonist TTA-P2 completely eliminated PF. In addition, replacing Ca^2+^ with Ba^2+^ abolished PF. These findings suggest that the Ca^2+^ increases required for TRPM4 activation are mediated by influx through T-type Ca^2+^ channels although we cannot rule out a contribution of R-type Ca^2+^ channels (Zaman et al., 2011) or release from internal stores (Neyer et al., 2016). Ca_V_3.3 channels mediating I_T_ are primarily expressed in the distal dendrites of TRN neurons and their activation during burst firing results in large dendritic but more modest somatic Ca^2+^ signals (Crandall et al., 2010; Astori et al., 2011). This would suggest a dendritic location of TRPM4. In this scenario, membrane depolarizations necessary for TRPM4 activation would likely be mediated by synaptically-generated T-type dependent Ca^2+^ spikes, which in contrast to fast Na^+^ action potentials, propagate effectively into TRN dendrites (Crandall et al., 2010; Connelly et al., 2015).

We found that in the presence of apamin, PF could be observed in most recordings, suggesting that TRPM4 conductances are expressed in the large majority of TRN neurons. Thus, SK channels control both the initiation and duration of PF, likely due to the close functional association with T-type Ca^2+^ channels (Cueni et al., 2008). Similarly, we found that the synaptic activation of M2 muscarinic receptors curtailed the duration of PF. It is therefore likely that under physiological conditions, TRPM4 activation and the generation of PF in a given neuron does not occur in all-or-none manner, but instead is constantly regulated by multiple synaptic and intrinsic processes. For example, an increased likelihood and duration of PF might be promoted by mechanisms that increase T-type mediated Ca^2+^ increases or directly facilitate TRPM4 activation, and curtailed by modulatory synaptic systems that reduce TRN excitability via opening of specific K conductances.

We want to emphasize that our findings do not provide evidence for a well-defined cell type that displays PF. Previous studies have used molecular markers (Clemente-Perez et al., 2017) or firing patterns (Lee et al., 2007) to delineate TRN neuronal subtypes that differ in the expression of T-type Ca^2+^ channels, which might be an important factor determining the incidence of PF. However, it is likely that a wide range of factors including the spatial arrangement and density of postsynaptic receptors, intrinsic conductances, and intracellular Ca^2+^ signaling mechanisms interact to control PF generation in a given cell.

### Comparison to previous studies

Plateau potentials and PF have been observed throughout the brain, though in most cases there is little consensus on the underlying mechanisms (Zylberberg and Strowbridge, 2017). In the thalamus, Kim and McCormick (1998) reported plateau potentials in a fraction of neurons of the perigeniculate sector of the ferret TRN, with very similar properties as described here. However, they concluded that plateau potentials were mediated by I_NaP_, since they were fully blocked by TTX. The same conclusion was reached in an *in vivo* study of TRN in anesthetized cats, which demonstrated block of persistent firing using intracellular dialysis with QX-314 (Fuentealba et al., 2005). Here we find that in apamin a significant number of TRN neurons shows plateau potentials even in the presence of TTX. It is possible that under physiological conditions transient or persistent forms of voltage-gated I_Na_ indirectly aid PF by increasing activation of TRPM4 and T-type Ca^2+^ channels.

For both TRN and thalamic relay neurons, group I mGluR activation can trigger rhythmic forms of PF, mediated by the closure of a K^+^ leak conductance, allowing the generation of a long-lasting inward current mediated by the non-inactivating portion of T-type Ca^2+^ channel conductance (Hughes et al., 2002; Blethyn et al., 2006). Our present findings differ from these results in four important aspects: First, PF did not require mGluR activation and the resulting closure of K^+^ leak conductances. Second, in the absence of mGluR activation, PF did not occur in a cell-intrinsic manner but required fast synaptic inputs. Under our experimental conditions, both cholinergic inputs and bursts of glutamatergic thalamic inputs triggered PF, likely by generating global Ca^2+^ spikes (Connelly et al., 2015). Third, while T-type Ca^2+^ channels were important for the initiation of PF, our results are inconsistent with a critical role of I_T_ in generating long-lasting currents that lead to plateau potentials. We found that plateau potentials reached levels of ∼-45 mV, where T-type Ca^2+^ conductances at steady-state experience near full inactivation. Furthermore, the lack of PF in the presence of Ba^2+^, which should allow for the generation of a I_T_-mediated window current (Huguenard and Prince, 1992) suggests that T-type Ca^2+^ currents alone are unlikely to mediate PF. Fourth, using the potent and selective antagonist CBA (Ozhatil et al., 2018), we showed that I_CAN_ was essential for the generation of plateau potentials and was mediated by TRPM4. It should be stated that the majority of TRP antagonists employed in previous studies have a number of non-specific effects, so the validation of CBA as a selective TRPM4 antagonist under our experimental conditions will require independent confirmation.

### Mechanisms of slow thalamic oscillations

Accumulating evidence indicates that both cortex and thalamus can generate oscillations in isolation, suggesting that rhythmic activity in the intact thalamocortical system results from the complex interplay of multiple distinct oscillators (Crunelli and Hughes, 2010). While there has been extensive research on the cortical mechanisms mediating slow rhythms, the nature of thalamic pacemakers are less well understood. Previous work has shown that both TRN and relay neurons are capable of generating cell-intrinsic rhythms in the 1 Hz range, but only under conditions of postsynaptic mGluR activation (Hughes et al., 2002; Blethyn et al., 2006). Under physiological conditions, this would require sustained corticothalamic activity, indicating that both thalamic cell types act as conditional oscillators. Our findings derived from adult thalamic slices highlight an alternative mechanism and suggest that networks of TRN and VB cells can generate robust and long-lasting slow oscillatory activity, mediated by powerful bidirectional synaptic connectivity (Gentet and Ulrich, 2003; Pinault, 2004; Pita-Almenar et al., 2014). We observed an inverse relationship between TRN firing duration and oscillatory frequency. It is therefore tempting to suggest that changes in TRN activity as seen during PF acts to reduce the frequency of thalamic oscillations, via recruitment of GABA_B_Rs (Kim et al., 1997) or extrasynaptic high-affinity GABA_A_Rs (Herd et al., 2013), thereby changing the latency to rebound burst generation (Schofield et al., 2009).

Our network data suggest that robust thalamic rhythmicity can occur in the absence of precise synchrony, highlighting that both processes can occur independently. While thalamic oscillatory activity is commonly referred to as synchronous (Fogerson and Huguenard, 2016), there have been few studies that have directly examined the degree of precise thalamic synchronization during distinct behavioral states. Our current data showing a lack of precise synchrony in local thalamic circuits confirm previous findings (Pita-Almenar et al., 2014) and indicate that the variability in TRN firing might act as an additional factor in preventing synchrony in isolated thalamic networks. As has been suggested (Contreras and Steriade, 1996), this would argue that synchronization of thalamic networks is primarily regulated by afferent inputs, particularly from cortex.

## Acknowledgments

Supported by funds from the Harry S. and Isabel C. Cameron Foundation (J.O.), the Neurodegeneration Consortium and the Belfer Family Foundation (J.C. and M.B.), NIH NS085171, NS086965 (J.C.), and NIH NS077989 (M.B.). We thank Michael Zhu and Oleh Pochynyuk for donating mouse models, John Magnotti for statistical advice, and Rajan Dasgupta and Fabricio Do Monte for helpful discussions.

